# Computational tools to detect signatures of mutational processes in DNA from tumours: a review and empirical comparison of performance

**DOI:** 10.1101/483982

**Authors:** Omichessan Hanane, Severi Gianluca, Perduca Vittorio

**Affiliations:** CESP (UMR INSERM 1018), Université Paris-Saclay, UPSud, UVSQ, Villejuif Cedex, F-94805, France; Gustave Roussy, Villejuif, F-94805, France; Cancer Epidemiology Centre, Cancer Council Victoria, and Centre for Epidemiology and Biostatistics, Melbourne School for Population and Global Health, The University of Melbourne, Australia; Laboratoire de Mathématiques Appliquées – MAP5 (UMR CNRS 8145), Université Paris Descartes, 75006, Paris, France

**Keywords:** sequencing, cancer, mutational signatures, decomposition algorithms, probabilistic models, simulation study

## Abstract

Mutational signatures refer to patterns in the occurrence of somatic mutations that reflect underlying mutational processes. To date, after the analysis of tens of thousands of exomes and genomes from about 40 different cancers types, tens of mutational signatures characterized by a unique probability profile across the 96 trinucleotide-based mutation types have been identified, validated and catalogued. At the same time, several concurrent methods have been developed for either the quantification of the contribution of catalogued signatures in a given cancer sequence or the identification of new signatures from a sample of cancer sequences. A review of existing computational tools has been recently published to guide researchers and practitioners through their mutational signature analyses, but other tools have been introduced since its publication and, a systematic evaluation and comparison of the performance of such tools is still lacking. In order to fill this gap, we have carried out an empirical evaluation of the main packages available to date, using both real and simulated data.

## Introduction

It is well-known that cancer originates from a DNA modification in a single normal cell that is then propagated through cell divisions and that accumulates with further DNA modifications finally leading to abnormal, cancerous cells [1]. Such changes that include Single Nucleotide Variants (SNVs), insertions or deletions, Copy Number Variation (CNVs) and chromosomal aberrations, have been termed as “somatic mutations” to distinguish them with those inherited and transmitted from parents (germline mutations). With the development and the improvement of sequencing technologies collectively referred to as *High-Throughput Sequencing (HTS)* and the availability of cancer exome and genome data from most human cancers, much has been learnt about these mutations and the set of genes (oncogenes and tumour suppressor genes) that operate in human cancers. Beyond the relatively small number of “driver” mutations, those that confer a selective advantage for tumour development and progression, the vast majority of mutations, so-called “passenger” mutations, are “neutral” that means they do not affect cancer cells’ survival. Together, passenger and driver mutations constitute a record of all cumulative DNA damage and repair activities occurred during the cellular lineage of the cancer cell [2].

Among all types of somatic mutations, a particular focus has been placed on single base substitutions that have been classified in six types according to the mutated pyrimidine base (C or T) in a strand-symmetric model of mutation. Such 6 substitutions (C>A, C>G, C>T, T>A, T>C and T>G) may be further classified in different types when considering the sequence pattern in which they are located (sequence context). For practical reasons, the sequence context is typically defined using the 5’ and 3’ bases proximal to the mutated base, that results in substitutions being classified in 96 types (6 *\* 4 * 4*). In this paper we adopt this convention and use “mutations” to refer to these 96 types of contextualized substitutions.

It has been hypothesised that specific patterns of mutations are the results of the action of particular underlying mutational process. To identify such patterns, computational models, such as matrix decomposition algorithms or probabilistic models, have been developed. The first of such methods was published in 2013 by Alexandrov and colleagues [2,3], and, as for most of all the other models that followed, is based on the idea that a mutational signature can be seen as a probability distribution of the 96 types of mutations. Mutational signatures contribute to the total mutational burden of a cancer genome, the so called mutational “catalogue” or “spectrum”.

Such original methodological framework, applied to tens of thousands of genomes and exomes from 40 different cancers types from large data repositories such as TCGA (The Cancer Genome Atlas), has led to the identification of 30 mutational signatures characterized by a unique probability profile across the 96 mutation types. These validated mutational signatures are listed in a repertory on the COSMIC (Catalogue Of Somatic Mutations In Cancer) website [3] and have been widely used as references. More recently, Alexandrov et al. have introduced an updated set of signatures identified from an even larger collection of both exome and whole-genome sequences (including the sequences from the PCAWG project) using two different methods (a new version of the original framework and a Bayesian alternative) [4]. The new repertory includes 49 new mutational signatures based on single base substitutions as in COSMIC, and also mutational signatures built in the context of other types of mutations such as double base substitutions (11 signatures), clustered based substitutions (4 signatures) and small insertions and deletions (17 signatures).

After the introduction of the original framework, several other mathematical methods and computational tools have been proposed to detect mutational signatures and estimate their contribution to a given catalogue. These methods can be grouped in two categories with different goals. The first class of methods aims to discover novel signatures while the second class aims to detect the known and validated mutational signatures in the mutational catalogue of a given sample. The approaches used in the first class are referred to as “*de novo*” (or “signature extraction”) while those in the second class as “refitting” (or “signature fitting”). All methods have been implemented in open source tools, mainly R packages, but some of them are available through command line, the Galaxy project or a web interface.

Signatures identified with de novo methods can be compared to reference signatures (for instance those listed in COSMIC) through measures such as cosine [5] or bootstrapped cosine similarity [6], which is a distance metric between two non-zero vectors. In this step of the analysis, extracted signatures are matched to the most similar reference signature, provided that their similarity is greater than a fixed threshold.

Recently, Baez-Ortega and Gori published a paper that discussed the mathematical models and computational techniques for mutational signature analysis [7]. However, to our knowledge no systematic evaluation of the performance of the existing methods has been conducted and the issue of the choice of an appropriate cosine similarity threshold when matching a newly extracted signature to the most similar counterpart in a reference set has not been addressed yet.

The present work has been designed to use simulated and real-world cancer sequences in order to evaluate and compare for the first time the methods available to date, and assess their ability to accurately detect the underlying mutational signatures in mutational catalogues.

## Mathematical definition of mutational catalogues and signatures

In this section we briefly review the mathematical framework and notations based on the original model developed by Alexandrov and colleagues [2]. The data structure that is used as input by most current methods is the *mutational catalogue,* or *spectrum,* of a genome (or exome), with the counts for each of the 96 different mutation *features,* or *types.* In most approaches, the notion of mutation type refers to base substitutions within the trinucleotide context described above.

The mutational catalogue of a genome (or exome) *g* is defined as the vector 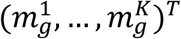, where each 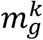 is the number of mutations of type *k* found in the genome and *K*, the number of possible mutation types, is equal to 96. Note that in this setting, information about mutation locations in the sequence is lost. The catalogue is built by comparing the sequence to a reference sequence in order to detect mutations and then by simply counting the occurrences of each tyspe. The reference sequence can either be a standard reference (e.g. the assembly GRCh38) or a sequence from a “normal” tissue from the same individual (e.g. DNA from blood or from normal tissue surrounding tumours when available). For the purposes of the present work we will use the generic term “samples” for both genomes and exomes as the concepts and models used may be applied to both.

The basic idea underlying all computational models proposed is that the mutational catalogue of a sample results from the combination of all the mutational processes operative during lifetime, and therefore it can be seen as the weighted superposition of simpler mutational *signatures,* each corresponding to a specific process. The weight is larger if the process has a larger role in the final catalogue of mutations: for example, for exposures to mutagens that last longer and/or are more intense.

Formally, the signature of a mutational process *n* is a vector 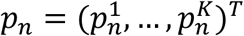, where each 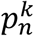 represents the probability that the mutational process will induce a mutation of type *k*. In other words, 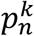 is the expected relative frequency of type *k* mutations in genomes exposed to *n*. Note that 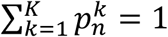 and 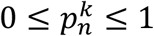 for all *k*.

The intensity of the exposure to a mutational process *n* in a sample *g* is measured by the number of mutations 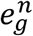 in *g* that are due to *n*. For this reason, 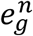 is referred to as the “exposure” of *g* to *n*. The expected number of mutations of type *k* due to the process *n* in sample *g* is therefore 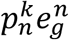. If sample *g* has been exposed to *N* mutational processes, then the total number of mutations of type *k* is

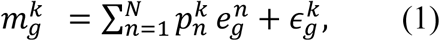

where 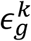 is an error term reflecting sampling variability and non-systematic errors in sequencing or subsequent analysis.

Matrix notation is effectively used when dealing with several samples and signatures. In this situation, the collection of *G* samples is represented by the *K* × *G* matrix, with catalogues in columns:

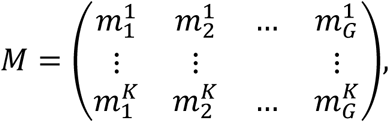

the *N* signatures are represented by the *K* × *N* matrix

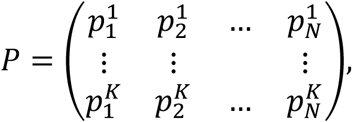

and the exposures by the *N* × *G* matrix

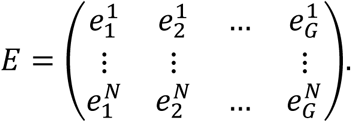

Equation (1) then becomes

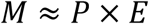

where we omitted the error term.

### *de novo* extraction vs. refitting

*de novo* signature extraction methods aim at estimating *P* and *E* given *M*. Non-negative matrix factorization (NMF) is an appealing solution to this unsupervised learning problem, because, by definition, all involved matrices are non-negative [8]. NMF identifies two matrices *P* and *E* that minimize the distance between *M* and *P* × *E*. In particular, NMF finds an approximated solution to the non-convex optimization problem

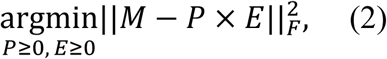

where the Frobenius matrix norm of the error term is considered. NMF requires the number of signatures *N*, an unknown parameter, to be predefined. An approach for selecting this parameter consists in obtaining a factorization of *M* for several of its values and then choosing the best *N* with respect to some performance measure such as the reconstruction error or the overall reproducibility. NMF is at the core of the Welcome Trust Sanger Institute (WTSI) Mutational Signature Framework, the first published method for signature extraction [2]. An alternative to numerical approaches based on NMF is given by statistical modelling and algorithms. With these latter approaches, the number of mutations of a given type can be modelled by a Poisson distribution

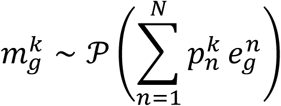

where mutational processes are assumed to be mutually independent. In order to estimate *E* and *P*, it has been proposed to consider *E* as latent data and *P* as a matrix of unknown parameters and to apply an expectation-maximization algorithm [9] or use Bayesian approaches [10–12]. One important advantage of statistical approaches is the availability of model selection techniques for the choice of *N*. We refer to the recent review [7] for a comprehensive overview of the mathematical and computational aspects of existing methods, including alternative techniques not mentioned above.

With respect to de novo extraction of signatures, the refitting approaches consider that the signatures *P* are known and the goal is to estimate *E* given *M* and *P*. Refitting can be done for individual mutational catalogues (i.e. individual samples) and, from a linear algebra perspective, can be seen as the problem of projecting a catalogue living in the *K*-dimensional vector space (the space spanned by all mutation types) onto its subset of all linear combinations of the given mutational signatures with non-negative coefficients (the cone spanned by the given signatures).

A current practice consists in first performing a de novo extraction of signatures followed by a comparison of the newly identified signatures with the reference signatures (e.g. COSMIC signatures) by means of a similarity score, typically cosine similarity ranging from 0 (completely different) to 1 (identical) [6,13]. Finally, a “novel” signature is considered to reflect a specific reference signature if the similarity is larger than a fixed cut-off. If similarity is observed with more than one reference signature, the one with the largest value of similarity is chosen, S1 Fig. This assignment step crucially depends on the choice of the cut-off *h* that has been so far inconsistent in the literature with some studies using a value of 0.75 [14] whereas others 0.80 [15,16]. Another difficulty is that different signatures might have very close cosine similarity, as it happens also between COSMIC signatures, S2 Fig, so that a unique assignment is not always possible.

## Overview of the available tools

An similar number of *de novo* and refitting methods exist and all of them are available as open source tools, mainly as R packages, or web interfaces (Table 1). The typical input of the tools is a file including the mutations counts but some tools derive the mutation counts from ad-hoc input files that may include for each individual a list of mutated bases, their position within the genome and the corresponding bases from a reference genome. The typical format of such input files is MAF, VCF or less common formats such as MPF and MFVF. For biologists or those who are not familiar with programming, a set of tools were also developed and provided with user-friendly interfaces (see also Table 1). Some tools include additional features such as the possibility to search for specific patterns of mutations (e.g. APOBEC-related mutations [17]) and differentially expressed genes [5].

| Software | Available platform/<br>model | Input files | Additional features |
| --- | --- | --- | --- |
| *de novo* approaches |  |  |  |
| WTSI | MATLAB/ NMF |  | Original framework |
| EMu [9]<br>https://github.com/andrej-fischer/EMu | Command line/EM algorithm | - Mutation counts file<br>- With respect to other tools, the counts file is transposed (the rows correspond to the samples) | - Opportunity matrix<br>- Selection of the optimal number of signatures |
| SomaticSignatures [17]<br>https://bioconductor.org/packages/release/bioc/html/SomaticSignatures.html | R/NMF and PCA | Variant Call Format | - Group-wise comparisons<br>- Genomic visualization<br>- Hierarchical clustering |
| pmsignature [18]<br>https://github.com/friend1ws/pmsignature | R/mixed-membership model | - Mutation Position Format<br>- Mutation Feature Vector Format | - Reduction of complexity<br>- Mutation types defined by one or two flanking bases<br>- Selection of the optimal number of signatures<br>- Transcriptional strand bias<br>- Background signature |
| bayesNMF [11,19,20]<br>https://github.com/jburos/bayesNMF<br>https://software.broadinstitute.org/cancer/cga/msp | R/Bayesian NMF | Mutation counts file | - Selection of the optimal number of signatures |
| signeR [10]<br>https://bioconductor.org/packages/release/bioc/html/signer.html | R/Bayesian NMF | Variant Call Format | - Opportunity matrix<br>- Selection of the optimal number of signatures<br>- Group-wise comparison (differential analysis) |
| mutSignatures [21]<br>https://cran.r-project.org/web/packages/mutSignatures/index.html | R/NMF | Mutation counts file | - R-based implementation of [2] |
| Maftools [5]<br>https://bioconductor.org/packages/release/bioc/html/maftools.html | R-Bioconductor /NMF | - Mutation Annotation<br>- Format | - Genomic visualization<br>- Cosine similarity<br>- Selection of the optimal number of signatures<br>- Group-wise comparisons (differential analysis)<br>- APOBEC enrichment analysis |
| Helmsman [22]<br>https://github.com/carjed/helmsman | Python/ NMF and PCA | - Variant Call Format<br>- Mutation Annotation Format | - Able to run in parallel and designed for large datasets<br>- Connection to external packages (in R)<br>- may generate mutational catalogues from sequence data |
| SigAnalyzer [23] | R/ Bayesian NMF | Mutation counts file | - Automatic selection of the optimal number of signatures<br>- Sparse signature profiles and contributions |
| SigProfiler [2,23] | Matlab/ NMF | Mutation counts file | - Further development of the original framework<br>- Two steps: 1) extraction of a minimal set of signatures, 2) estimation of their contributions to individual samples |
| SparseSignatures [24]<br>https://bioconductor.org/packages/release/bioc/html/SparseSignatures.html | R/ NMF with Lasso-penalized cost function | Mutation counts file | - Integration of DNA replication error signature<br>- Sparse signature matrix<br>- Number of signatures estimated with cross-validations<br>- Scalable to large datasets |
| **Refitting approaches** |  |  |  |
| deconstructSigs [25]<br>https://github.com/raerose01/deconstructSigs | R/linear regression | Mutation counts file | - Opportunity matrix |
| QPsig [26]<br>https://f1000researchdata.s3.amazonaws.com/supplementary/8918/0d25c07c-16ba-4b14-91e7-71749dcbdd5.pdf | R/quadratic programming | Mutation counts file |  |
| SignatureEstimation [27]<br>https://www.ncbi.nlm.nih.gov/CBBresearch/Przytycka/index.cgi/#signatureestimation | R/quadratic programming and simulated alienation | Mutation counts file |  |
| MutationalPatterns [28]<br>http://bioconductor.org/packages/release/bioc/html/MutationalPatterns.html | R/Non-Negative Least Squares | Mutation counts file | - Also de novo identification<br>- Cosine similarity comparison<br>- Strand bias analyses<br>- Enrichment and depletion |
| YAPSA [29]<br>http://bioconductor.org/packages/release/bioc/html/YAPSA.html | R/Linear Combination Decomposition | Mutation counts file | - Cut-off for normalized exposure<br>- Enrichment and depletion |
| decompTumor2Sig [30]<br>https://github.com/rmpiro/decompTumor2Sig | R/quadratic programming | - Variant Call Format<br>- Mutation Position Format<br>- Mutation Feature Vector Format | - Converts a set of "Alexandrov's signatures" [13] to "Shiraishi's signatures" [18]<br>- Decomposes a mutational catalogue in "Shiraishi's signatures" |
| MutationalCone [See Supporting information] | R/cone projection | Mutation counts file | - Fast in comparison to others refitting tools |
| Sigfit [12]<br>https://github.com/kgori/sigfit | R/ Bayesian NMF | Mutation counts file | - Provides a new model for combining de novo and refitting approaches<br>- Possible application to indel or rearrangement count data<br>- Also implements Emu [9] model and allows conversion to genome-or exome- relative signatures |
| **Pipelines and web-interfaces** |  |  |  |
| Mutspec [31]<br>https://toolshed.g2.bx.psu.edu/repository/view_repository?id=f5c1f75e9fb33f8e | Galaxy pipeline/NMF | Variant Call Format | - de novo identification<br>- Includes MS analysis in mouse cancer |
| MutaGene [32]<br>https://www.ncbi.nlm.nih.gov/research/mutagene/ | Web-interface | TCGA and ICGC data | - Refitting and de novo identification<br>- Clustering of samples according to mutational profiles<br>- Identification of potential driver's mutations |
| mSignatureDB [6]<br>http://tardis.cgu.edu.tw/msignaturedb/ | Web-interface | - Variant Call Format<br>- Mutation Annotation Format<br>- TSV | - Refitting and de novo identification<br>- Bootstrapped cosine similarity<br>- Comparison with either hg19 or hg38 |
| Mutalisk [33]<br>http://mutalisk.org | Web-interface | Variant Call Format | - Refitting and de novo identification<br>- Transcriptional strand bias<br>- Localization of kaetegis<br>- Histones modifications<br>- Cosine similarity comparison |
| MuSiCa [34]<br>http://bioinfo.ciberehd.org:3838/MuSiCa/ | Web-interface | - Variant Call Format<br>- Mutation Annotation Format<br>- TSV<br>- Excel | - Refitting and de novo identification<br>- Cosine similarity<br>- Samples classification |

### *de novo* approaches

Most tools that have been developed to identify mutational signatures were based on decomposition algorithms including NMF or a Bayesian version of NMF [10,11]. The original method developed by Alexandrov et al. was based on NMF and was implemented in MATLAB [2] and is available also as an R package developed independently [21]. An updated and elaborated version named SigProfiler, was proposed recently for extracting a minimal set of signatures and estimating their contribution to individual samples [4]. The latter article also discusses an alternative method based on Bayesian NMF, called SignatureAnalyzer, that led to the identification of 49 reference signatures. Another tool that utilises NMF is Maftools that is one of the few *de novo* tools that allows systematic comparison with the 30 validated signatures in COSMIC by computing cosine similarity and assigning the identified signatures to the COSMIC one with the highest cosine similarity [5].

Other tools such as SomaticSignatures [17] or the recent Helmsman [22] allows the identification of mutational signatures through Principal Component Analysis (PCA) in addition to NMF. For the sake of our formal comparison of the tools performance, we have tested SomaticSignatures only with the NMF implementation because with PCA the factors are orthogonal and the values inside the matrix can potentially be null or negatives, which is biologically unrealistic. Developed in the Python language, Helmsman allows the rapid and efficient analysis of mutational signatures directly from large sequencing datasets with thousands of samples and millions of variants.

A promising recent method called SparseSignatures [24] proposes an improvement of the traditional NMF algorithm based on two innovations, namely the default incorporation of a background signature due to replication errors during cellular division and the enforcement of sparsity in identified signatures through a Lasso penalty. This latter feature allows the identification of signatures with well-differentiated profiles, thus reducing the risk of overfitting.

In addition to decomposition methods, an approach based on the Expectation Maximization (EM) algorithm has been proposed to infer the number of mutational processes operative in a mutational catalogue and their individual signatures. This approach is implemented in the EMu tool [9], where the underlying probabilistic model is based on the assumption that the input samples are independent and the number of mutational signatures is estimated using the Bayesian Information Criterion (BIC).

Another tool that uses a probabilistic model named mixed-membership model is pmsignature [18]. This tool utilises a flexible approach that at the same time reduces the number of estimated parameters and allows to modify key contextual parameters such as the number of flanking bases. The latter feature makes it possible to consider more-fine grained signatures and could potentially lead to new biological insights [23]. It is important to note that for the purpose of our comparison with the other tools, we set to one the number of flanking bases, and therefore to 96 mutation types.

EMu, SigneR and pmsignature (and the refitting tool deconstructSigs) have been designed to take into account the distribution of triplets in a reference exome or genome for example from a sequence of normal tissue in the same individual. This is done by “normalizing” the input mutational catalogues with respect to the distribution of triplets in the reference exome or genome using an “opportunity matrix”.

## Refitting with known mutational signatures

In addition to the identification of novel mutational signatures, we are often interested in evaluating whether a signature observed in an individual tumour is one of an established set of signatures (e.g. the COSMIC signatures). This task is performed by “refitting tools” that aim to search for the “best” combination of established signatures that explains the observed mutational catalogue by projecting the latter (i.e. mapping) into the multidimensional space of all non-negative linear combinations of the *N* established signatures.

The deconstructSigs [25] tool searches for the best linear combination of the established signatures through an iterative process based on multiple linear regression aimed at minimizing the distance between the linear combination of the signatures and the mutational catalogue. All the other tools minimise the distance through equivalent approaches based on quadratic programming [26,27,30], non-negative least square [28] linear combination decomposition [29] and simulated annealing [27].

### A faster implementation

We propose an alternative implementation of the decomposition performed in [28,29] based on a simple geometric framework. Finding the linear decomposition of the input catalogue *M* on a set of given signatures minimizing the distance can be seen as the problem of projecting *M* on the geometric cone whose edges are the reference signatures. We propose to solve this problem by applying the very efficient R package called coneproj [35]. More details about our algorithm, which we called MutationalCone, together with the R code implementing it, can be found in the Supporting information.

## Combining de novo and refitting procedures

Sigfit [12] is a recently introduced R package for Bayesian inference based on two alternative probabilistic models. The first of such models is a statistical formulation of classic NMF where signatures are the parameters of independent multinomial distributions and catalogues are sampled according to a mixture of such distributions with weights given by the exposures, while the second model is a Bayesian version of the EMu model. An interesting innovation of Sigfit is that it allows the fitting of given signatures and the extraction of undefined signatures in the same Bayesian process. As argued by the authors, this unique feature might be helpful in cases where the small sample of catalogues makes it difficult to try to identify new signatures or when the aim is to study the heterogeneity between the primary tumour and metastasis in terms of the signatures they show.

In this work we empirically evaluate the methods that have been already presented in a peer review published paper and for which an implementation in R is available. To this aim, we adopt the COSMIC set as reference for the analysis of simulated and real mutational catalogues because we evaluate tools that were developed at the time when COSMIC was the only available database of reference.

## Data and experimental settings

### Real cancers data from TCGA

In order to evaluate the performance of the available algorithms on real data, we used the exome sequences in The Cancer Genome Atlas (TCGA) repository (https://cancergenome.nih.gov/) for four cancer types: breast cancer, lung adenocarcinoma, B-cell lymphoma, and melanoma.

Mutation Annotation Format (MAF) files with the whole-exome somatic mutation datasets from these cohorts were downloaded from the portal gdc.cancer.gov on 6 March 2018. Data were annotated with MuSE [36] and the latest human reference genome (GRCh38). Mutational catalogues from these cohorts were obtained by counting the number of different mutation types using maftools [5]. The distribution of the number of mutations for each sample and separately for each cancer type is depicted in Fig 1.

**Fig 1.**
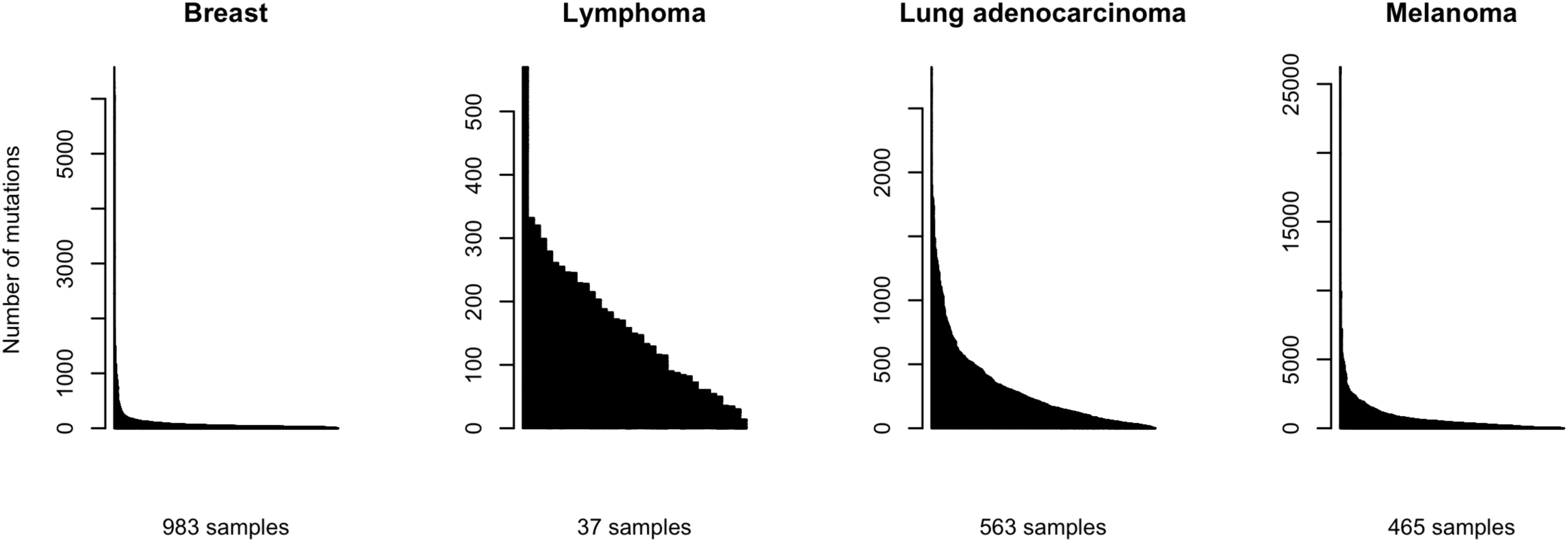
Barplot with the number of mutations in each sample in four TCGA cohorts. Each bar represents a sample, with the number of mutations shown in the y-axis.

According to the COSMIC website, 13 and 7 signatures have been found for breast cancer and lung adenocarcinoma respectively, 6 for B-cell lymphomas and 5 for melanoma.

## Simulated data

The first key assumption of our model of simulated mutational catalogues is that the number of mutations in a sample *g* that are induced by process *n* follows a zero-inflated Poisson (ZIP) distribution. According to this two-component mixture model, 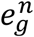 is either 0 with probability *π* or is sampled according to a Poisson distribution *𝒫*(*λ*) with probability 1 – *π*. Such a model depends on two parameters: the expectation *λ* of the Poisson component, and the probability *π* of extra zeros. The ZIP model allows for frequent zeroes and is therefore more suitable for modelling a heterogeneous situation where some samples are not exposed to a given mutational process 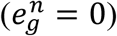 while some others are 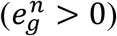. Realistically, the mutation counts due to process *n* in each of the *G* samples, 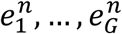, distributed according to a ZIP model where the expectation of the Poisson component is specific to *n*:

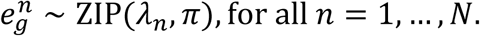

Note that the expected number of mutations in sample *g* due to process *n* is (1 – *π*)*λ*_*n*_. This flexibility given by process-specific average counts is the second important characteristics of our model and reflects the possibility that the mutagenic actions of different processes are intrinsically different with respect to their intensity. Obviously, it would have been possible to do one step further and allow for parameters *λ*_*n,g*_ specific to both processes and samples, thus representing the realistic situation in which the exposures of different samples to the same process have different duration (e.g. smokers/non-smokers). However, this would have resulted in too many parameters to tune, thus making it difficult to interpret the results of our simulation study. For the same reason we considered one fixed value of *π*.

The parameter *λ*_*n*_ depends on both the average total number *r* of mutations in a sample and the relative contribution of *n*. We therefore imposed the parameterization *λ*_*n*_(1 – *π*) = *q*_*n*_*r*, where *q*_*n*_ is the average proportion of mutations due to the process *n*.

When taking a unique value of *r*, this model produces realistic simulations even though it underrepresents extreme catalogues with very large or small total numbers of mutations (see S3 Fig (c)). While considering a specific value of *r* for each sample, or group of samples, would definitely make it possible to obtain a more realistic distribution of simulated catalogues (see S3 Fig (b)), such multidimensional parameter would complicate unnecessarily our empirical assessment of mutational signatures detection methods by introducing too many specifications. We therefore decided to consider a unique *r* for each set of simulations. This formulation allowed us to study empirically the performance of a given signature detection method as a function of the average number of mutations *r* while fixing the average proportion of mutations due to each mutational process *q*_*n*_, according to different profiles that mimic real cancer catalogues. Interestingly, the ZIP model appeared to be more appropriate to represent mutational catalogues than the pure Poisson model used in previous publications [9,10] (S3 Fig (d)).

Our simulation protocol was the following:

1. We chosed *N* signatures from the COSMIC database, thus obtaining the matrix *P*.
2. For each sample *g* and process *n* we sampled 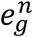 from a ZIP distribution with parameters *λ*_*n*_ = *q*_*n*_*r*/(1 – *π*) and *π* and obtained *E*. Here *q*_*n*_, *r* and *π* are fixed parameters to set.
3. We computed the product *P* × *E*. In order to obtain the final simulated catalogue *M*, we added some noise to the latter matrix by taking 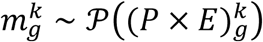.

We generated four alternative sets of simulated catalogues, referred to as Pofiles 1, 2, 3 and 4, each set mimicking a particular cancer: breast cancer, lung adenocarcinoma, B-cell lymphoma and melanoma. In order to do so, for each tumour type, we applied MutationalCone to the corresponding TCGA datasets and we calculated the mean contribution across all samples of each signature known to contribute to the specific cancer type *q*_*n*_. Signatures with *q*_*n*_ = 0 do not contibute to the final catalogue and were not in the matrix *P* S4 Fig depicts the resulting four sets of configurations (*q*_1_, ·, *q*_*N*_) used for the simulations. We fixed the frequency of structural null contributions to the catalogues to *π* = 0.6 in all simulations. This value was chosen because it leads to a limited number of hypermutated catalogues, as it is often encountered in practice. At last, we varied the number of *r* from as little as 10 to as much as 100,000 mutations. This allowed us to study the performance of methods on a large spectrum of catalogues: from a limited number of mutations as in exomes, to a very large number, as in whole cancer genome sequences.

For each of the four tumour types and for each value of *r*, we simulated a catalogue matrix with *G* samples. Fig 2 shows examples with *G* = 30 for three values of *r*.

**Fig 2.**
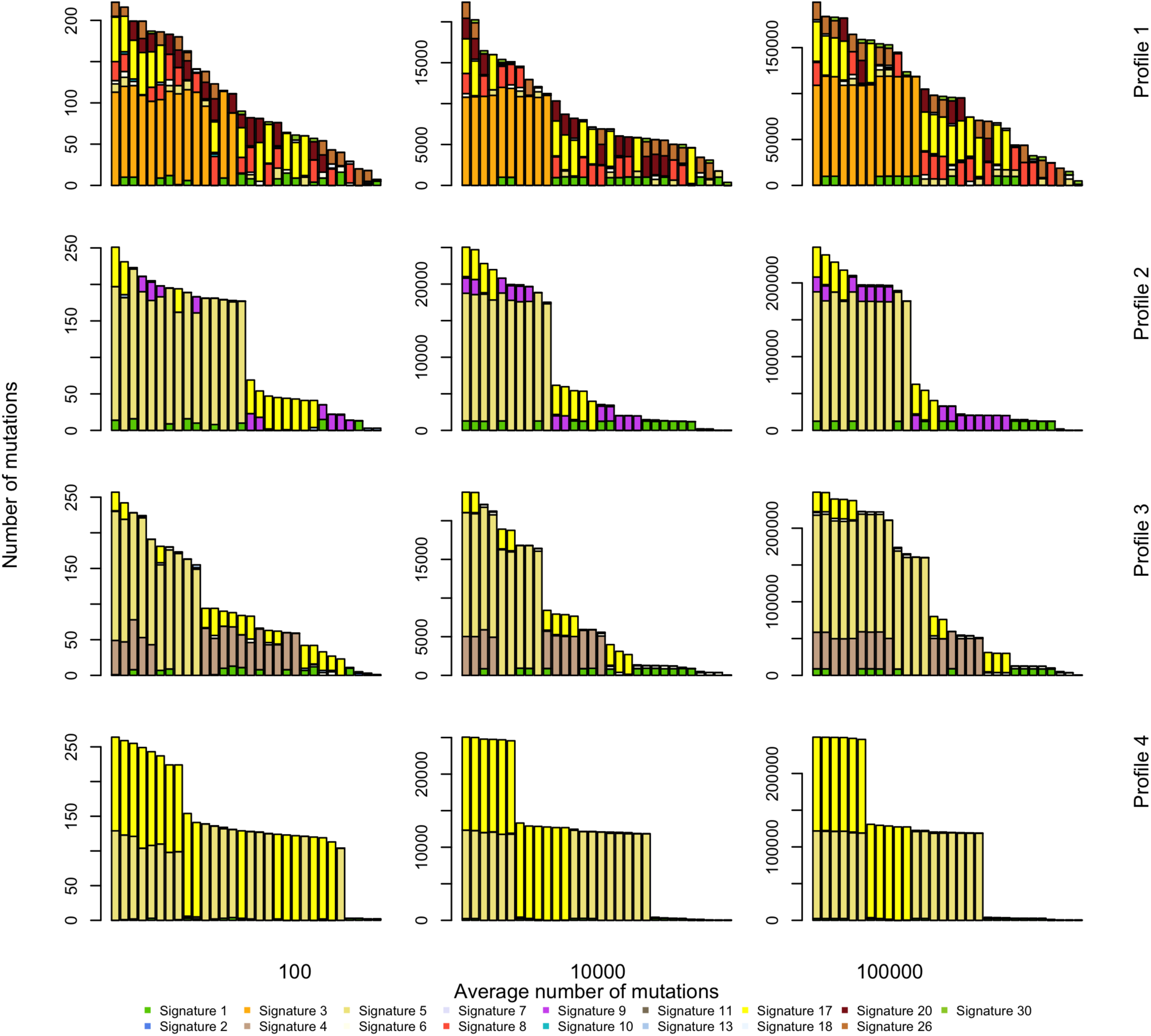
Sample of simulated catalogues. Catalogues are simulated according to the model described in the text. Tuning parameters are the average number of mutations, here *r* = 10^2^, 10^4^ and 10^5^ and the cancer type-specific average contribution of each signature (also see S4 Fig). Coloured bars indicate the contribution of each signature to the total mutational load of each sequence.

## Methods for measuring the performance of algorithms

All methods for identifying signatures find solutions to the minimization problem (2). A straightforward way to measure the accuracy of the reconstructed catalogue is, therefore, to calculate the Frobenius norm of the reconstruction error

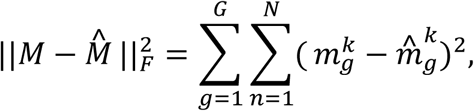

where 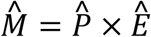 is the matrix of catalogues reconstructed from the estimated signature and exposure matrices. Some of the algorithms involve stochastic steps such as resampling and/or random draws of initial parameters. For these algorithms, one simple way to assess the robustness of the estimates is to look at the variability of the reconstruction error when the same catalogues are analyzed several times with the same algorithm.

In order to make decisions about whether an extracted signature is the same as validated signatures (e.g. COSMIC signatures) a cut-off for cosine similarity needs to be defined. We decided to apply six different cut-offs and consider as “new” all identified mutational signatures for which the maximal cosine similarity is lower than the cut-off value.

### Specificity and sensitivity for de novo extraction and assignment

In most applications, signature extraction is done in two steps: first, signatures are found using a de novo extraction tool and then for the extracted signatures a cosine similarity with each of the COSMIC signatures is calculated. In order to measure the performance of both these steps combined, we used simulated catalogues and computed false and true positive rates and false and true negative rates. In a simulated catalogue, the set of *true* signatures *p*_1_, ·, *p*_*N*_ that do contribute to the catalogue are known, thus allowing the comparison of the latter to the estimated signatures 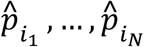. Note that, for the sake of simplicity, we explicitly look for the same number of signatures that we used to simulate the catalogues, not addressing questions about model selection performance.

Estimated signatures that belong to the set of “true signatures” are considered as true positives, while all “true signatures” that are not extracted count as false negatives. False positives are all estimated signatures that do not have a match in the set of “true signatures”. This can happen for two reasons: the estimated signature is assigned to a COSMIC signature not used to build the catalogue, or it is not sufficiently similar to any COSMIC signature. This last situation usually takes place when setting a very high cosine similarity threshold *h*. In this case, signatures that have maximal cosine similarity lower than the cutoff, will be termed as “new”. At last, true negatives are all COSMIC signatures not used for the simulation, nor estimated. From these four measures, we compute specificity (number of true negatives divided by the total number of negatives) and sensitivity (number of true positives divided by the total number of positives).

In our empirical study, for each simulation setting described in the Simulated data section (that is for each profile given by a choice of proportions (*q*_1_, ·, *q*_*N*_) and for a choice of total number of mutations *r*) we built 50 replicates, each made of a matrix of *G* samples. Then we extracted signatures from all replicates with a given tool. Importantly, we set the expected number of signatures to be the same as the number of signatures used in the simulation, thus not addressing the problem of model selection. Then we compared the extracted signatures to the COSMIC signatures using a cosine similarity threshold *h*. Finally, we computed specificity and sensitivity and obtained Monte-Carlo estimates based on the means over all replicates.

### Bias of refitting procedures

Because refitting algorithms assume that the matrix of signatures is known, a simple way to assess their performance is to look at the bias of the exposure estimates 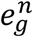. In order to do so, we simulated 50 replicates each consisting of one lymphoma-mimicking catalogue (Profile 2) with an average number of mutations set to *r* = 10^4^. Then, for each process *n*, we obtained Monte-Carlo estimates of the bias 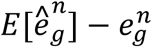 by averaging the differences 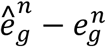 over all replicates. A global measure of performance that considers all exposure estimates is given by the mean squared error (MSE), that is the expected value of the loss function 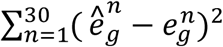. We obtained Monte-Carlo estimates of the mean squared error by averaging the loss function values across all 50 replicates and calculate asymptotic confidence intervals.

Analyses were performed using R 3.5.1 on a Linux Ubuntu (14.04) machine with 32 GB RAM and Intel Xeon E5-2630 v4 CPUs @ 2.2 GHz x 16.

## Comparisons of algorithm performance

### Performance of de novo tools

Fig 3 shows the distribution of the reconstruction error when a given computational tool is applied several times to the same real trinucleotide matrix. Reconstruction errors show limited variability due to stochastic steps in the algorithms and no variability whatsoever for EMu and Maftools. All methods under evaluation are roughly equivalent in terms of their ability to properly reconstruct the initial matrix of mutational catalogues. This is not surprising, given that all methods are meant to solve the optimization problem in (2). The error value also appears to be cancer-dependent. This is expected because cancers differ with regards to the total number of mutations and the number of operating mutational signatures, making the decomposition more or less difficult.

**Fig 3.**
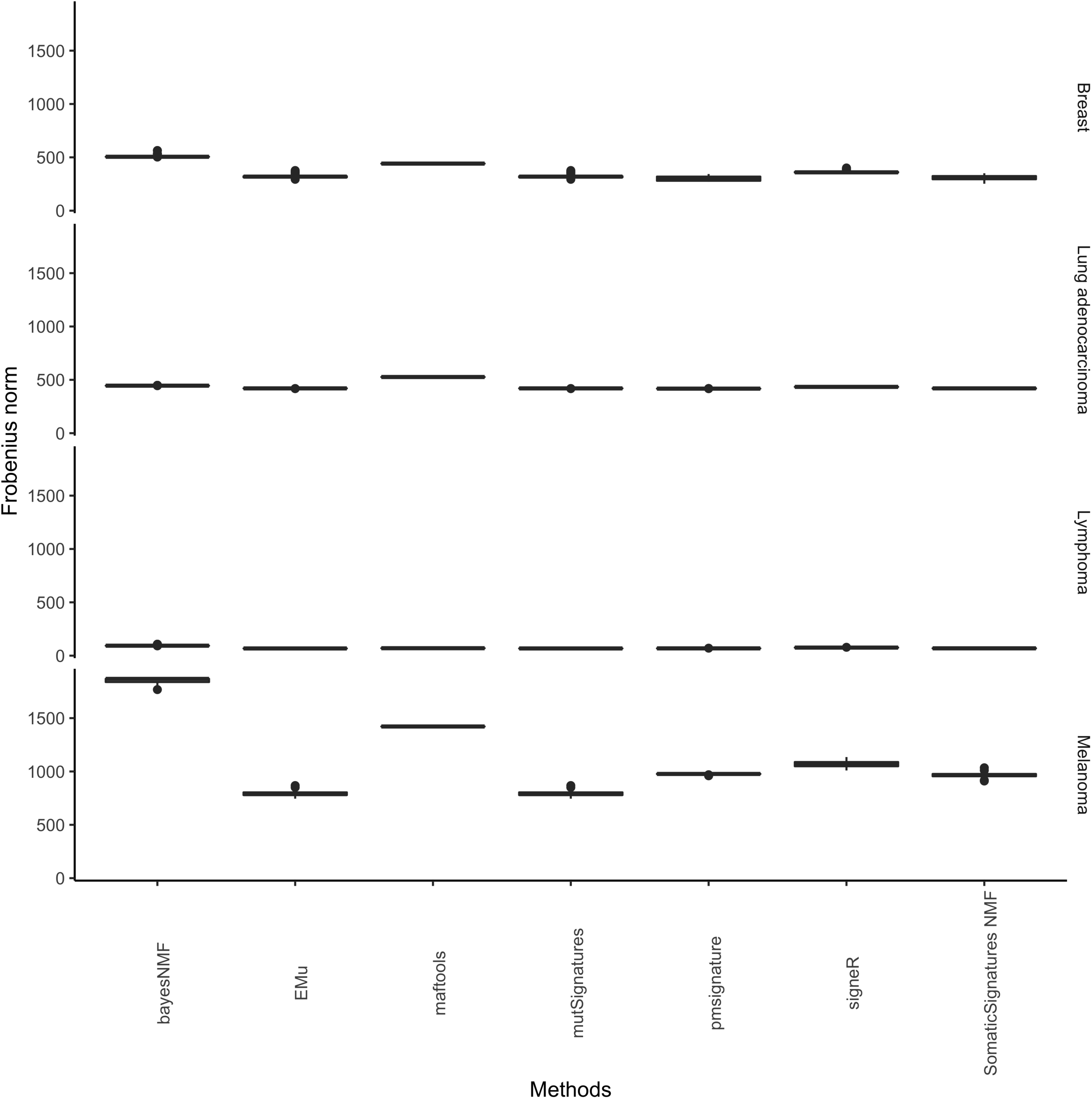
Reconstruction errors and their variability due to stochastic steps in the algorithms. Each program under evaluation is applied 50 times on the same matrix of real catalogues shown in Fig 1; boxplots represent the distribution of the squared Frobenius distance between the original catalogue and its reconstruction. Boxplots look like flat segments because of the scale of the y-axis.

We used realistic simulations to evaluate the performance of each method for de novo extraction followed by a classification step in which the extracted signatures are assigned to the most similar COSMIC signature.

Figs 4 and 5 show the specificity and sensitivity of such two-stage procedure as functions of the number of samples *G* in each catalogue and the cosine similarity cut-off *h*, while Figs 6 and 7 show the specificity and sensitivity as functions of the number of mutations in each catalogue and *h*.

**Fig 4.**
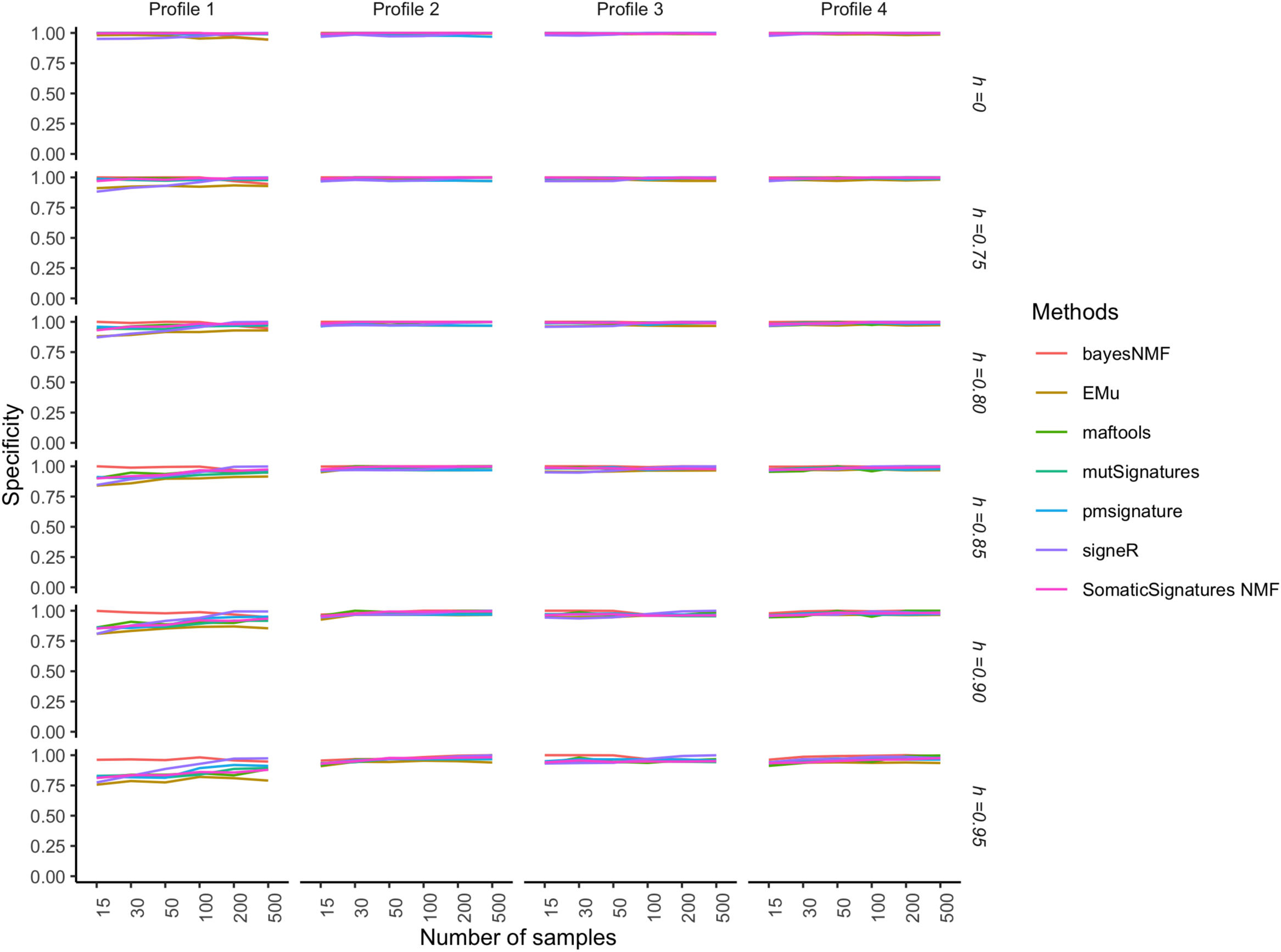
Performance of extraction methods and mapping on COSMIC signatures as the number of analysed catalogues and the cosine cut-off *h* vary. Specificity and sensitivity are estimated from 50 replicates each made of *G* genomes. The average number of mutations in each catalogue is *r* = 10,000. The model used to simulate realistic replicates according to the four Profiles and the estimation methods are described in the section Data and experimental settings.

**Fig 5.**
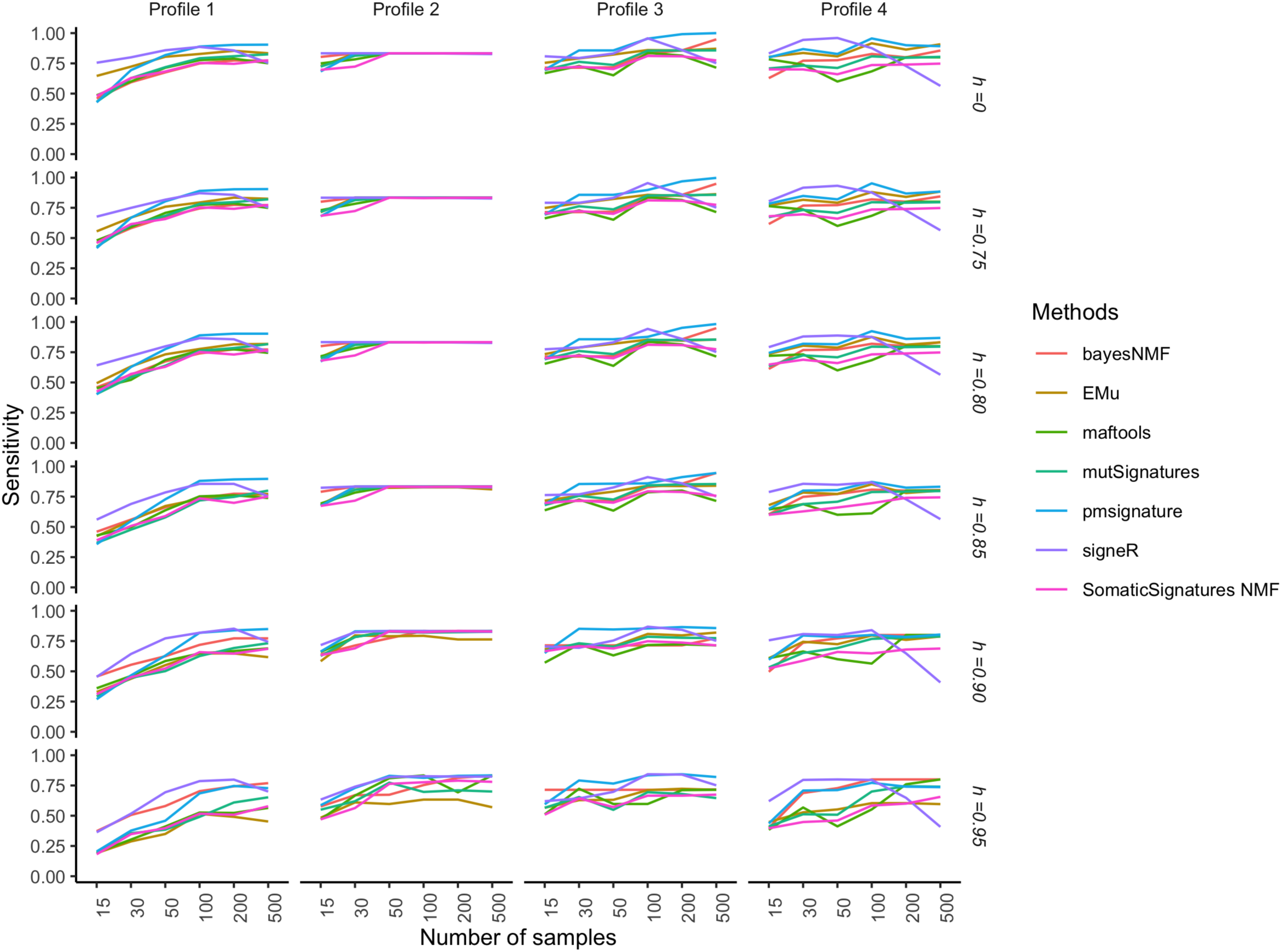
Performance of extraction methods and mapping on COSMIC signatures as the number of analysed catalogues and the cosine cut-off *h* vary. Specificity and sensitivity are estimated from 50 replicates each made of *G* genomes. The average number of mutations in each catalogue is *r* = 10,000. The model used to simulate realistic replicates according to the four Profiles and the estimation methods are described in the section Data and experimental settings.

**Fig 6.**
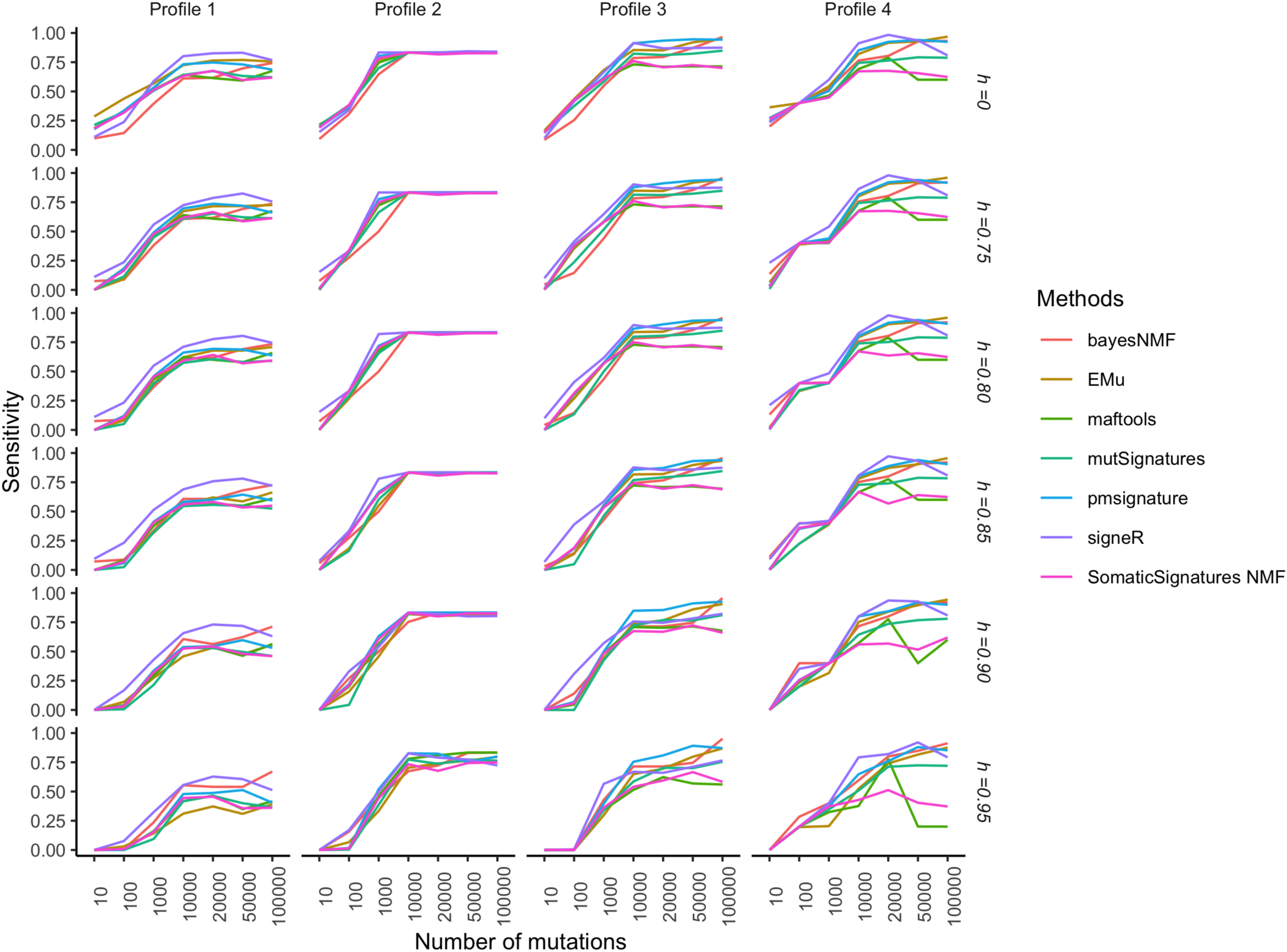
Performance of extraction methods and mapping on COSMIC signatures as the average number of mutations and the cosine cut-off *h* vary. Specificity and sensitivity are estimated from 50 replicates each made of *G* = 30 catalogues. The model used to simulate realistic replicates according to the four Profiles and the estimation methods are described in the section Data and experimental settings.

**Fig 7.**
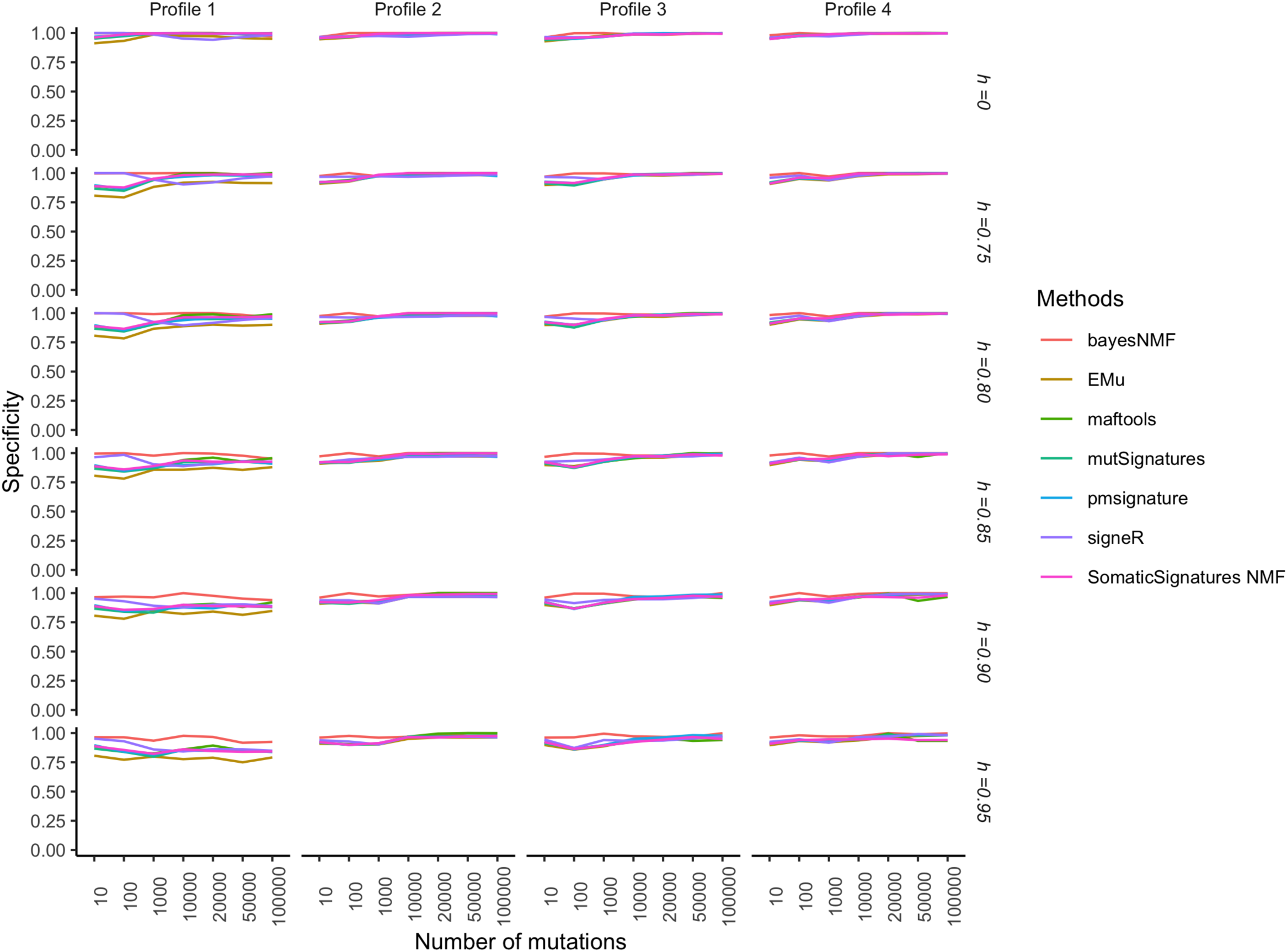
Performance of extraction methods and mapping on COSMIC signatures as the average number of mutations and the cosine cut-off *h* vary. Specificity and sensitivity are estimated from 50 replicates each made of *G* = 30 catalogues. The model used to simulate realistic replicates according to the four Profiles and the estimation methods are described in the section Data and experimental settings.

We do not see very large differences in the tools’ performance (specificity and sensitivity) between profiles with respect to the number of samples. For Profiles 2, 3 and 4 there is almost no gain in specificity and sensitivity when the sample size increases whereas for Profile 1, that is characterised by small contributions from several signatures (S4 Fig), sensitivity is poor for small samples sizes and increases with the sample size.

Sensitivity is poor when the average number of mutations is poor. Sensitivity increases with the average number of mutations with a large variability according to the cancer profile and method. In general, sensitivity is higher for profiles characterized by a signature with a stronger effect than other contributing signatures (e.g. Profile 2 vs. Profile 4). Specificity is less variable and generally high.

Specificity and sensitivity slightly deteriorate for higher cut-off values. This is expected because by setting a higher cut-off, the number of found signatures that are not similar enough to COSMIC signatures increases. Because these estimated signatures are considered as novel, they are false positives (that is found signatures not used for simulations), leading to a greater number of false positives and therefore to a lower specificity. Moreover, if the cut-off is too stringent, the number of false negatives will be high because some signatures used for the simulations are correctly found but do not score a high enough cosine similarity and therefore count as false negatives. This will make the resulting sensitivity low.

Fig 8 shows the running time when tools are applied to real lung datasets with a varying number of samples. While all methods show a fast-growing running time with increasing number of samples, SomaticSignatures and maftools are much faster than the others for more than 100 samples, making it possible to analyse large number of samples in few seconds. For example, for two hundred samples, the slowest method (SigneR), the running time is 913.72s while for the fastest (maftools), the value is 5.97s.

**Fig 8.**
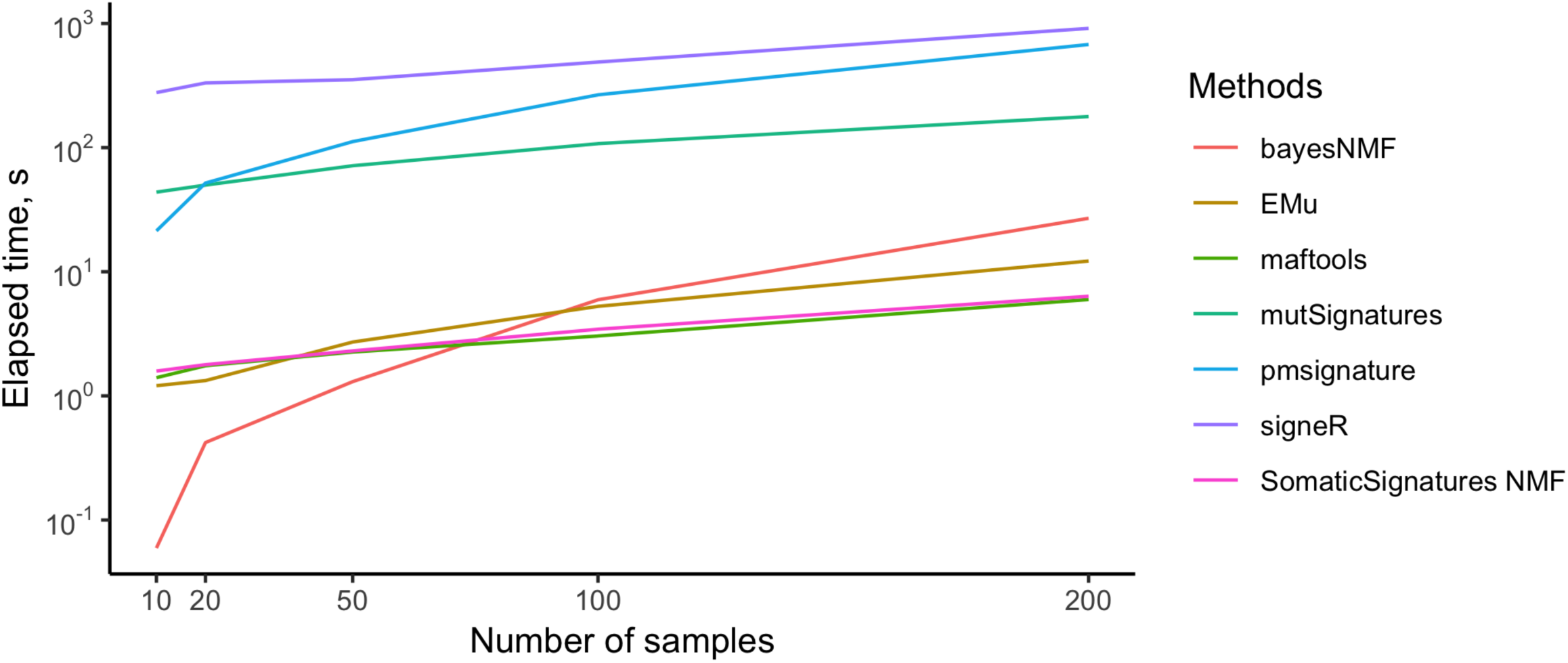
Running times of refitting tools. Methods were applied to subsets of the TCGA Lung cohort of different sizes.

### Performance of refitting tools

Fig 9 shows the distribution of the differences between the estimated and true contribution of all *n* signatures 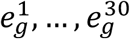 for the different refitting methods under evaluation. Sample catalogues were simulated mimicking Lung cancer profiles (Profile 3), with signatures 1,2,4,5,6, 13 and 17 actually contributing as shown in S4 Fig. All methods give almost identical results.

**Fig 9.**
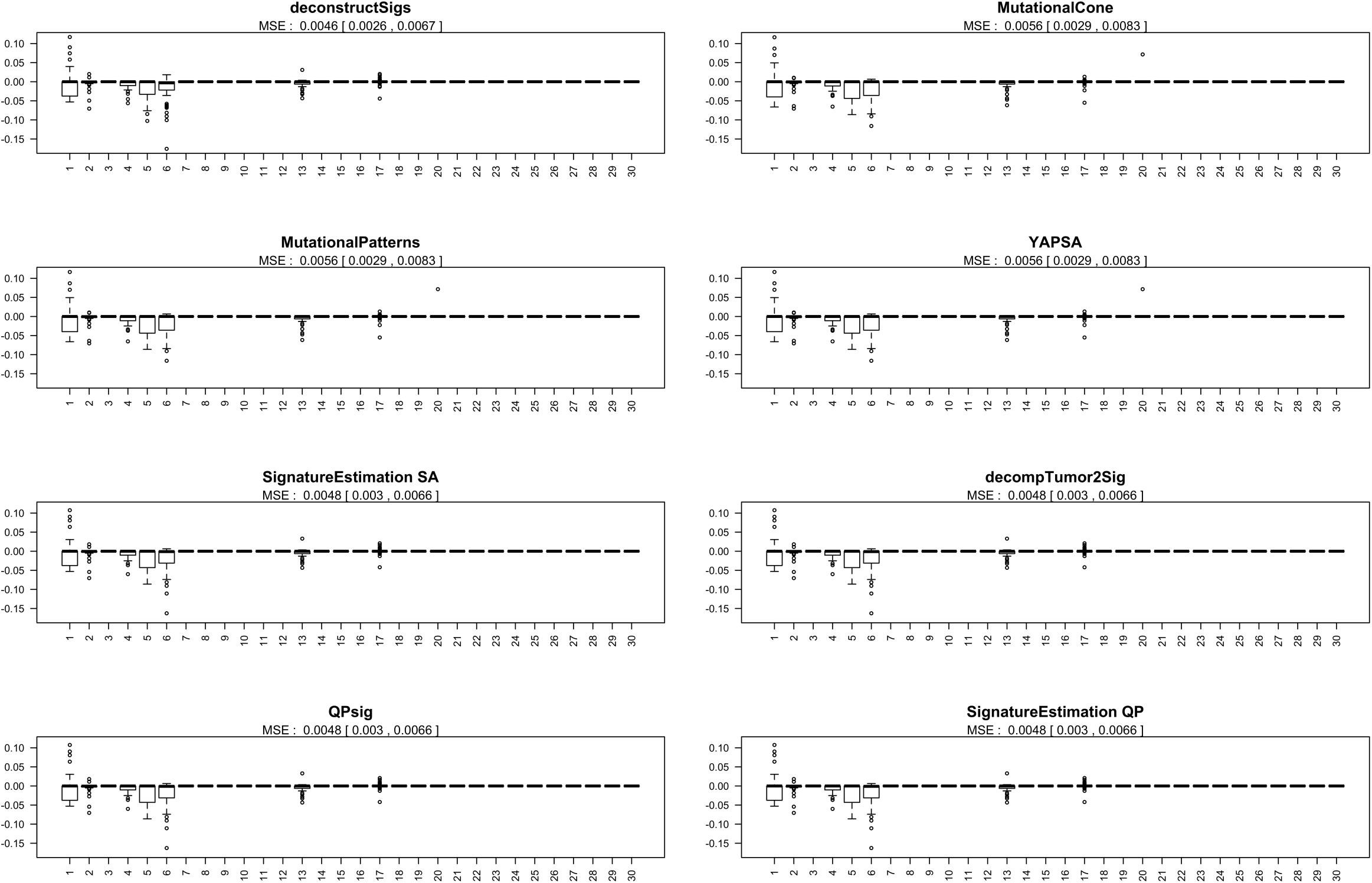
Bias of the estimates of each signature contribution for several refitting methods. For each signature, the bias estimates are obtained by averaging the exposure estimates across 50 samples. Mean square errors, together with 95% confidence intervals, are reported on the top of each plot. Simulations were done according to the model described in the Data and experimental settings section.

By comparison with the true exposure profile given in S4 Fig, it is clear that all refitting methods provide good estimates of the contributions of all but signatures 1,5 and, 6 and to a lesser extent signature 4. Moreover, all methods correctly estimate a zero contribution for signatures 3 and 16 even though these are very similar to signature 5, S2 Fig.

Interestingly, we note that signatures 2 and 13 (both attributed to APOBEC activity) are well identified by all methods. This finding is in line with previous claims about the stability of these two signatures.

In terms of running time, deconstructSigs and SignatureEstimation based on simulated annealing are more than two orders of magnitude slower than the other methods (Fig 10). All other methods run in a fraction of second. As expected, the running time increases linearly with the number of samples. MutationalCone, our custom implementation of the solution to the optimization problem solved by YAPSA and MutationalPatterns outperforms all other methods. The second fastest method is SignatureEstimation based on Quadratic Programing. As example, for two hundred samples, the execution time of deconstrucSigs is 86.148s and for MutationalCone is 0.028s.

**Fig 10.**
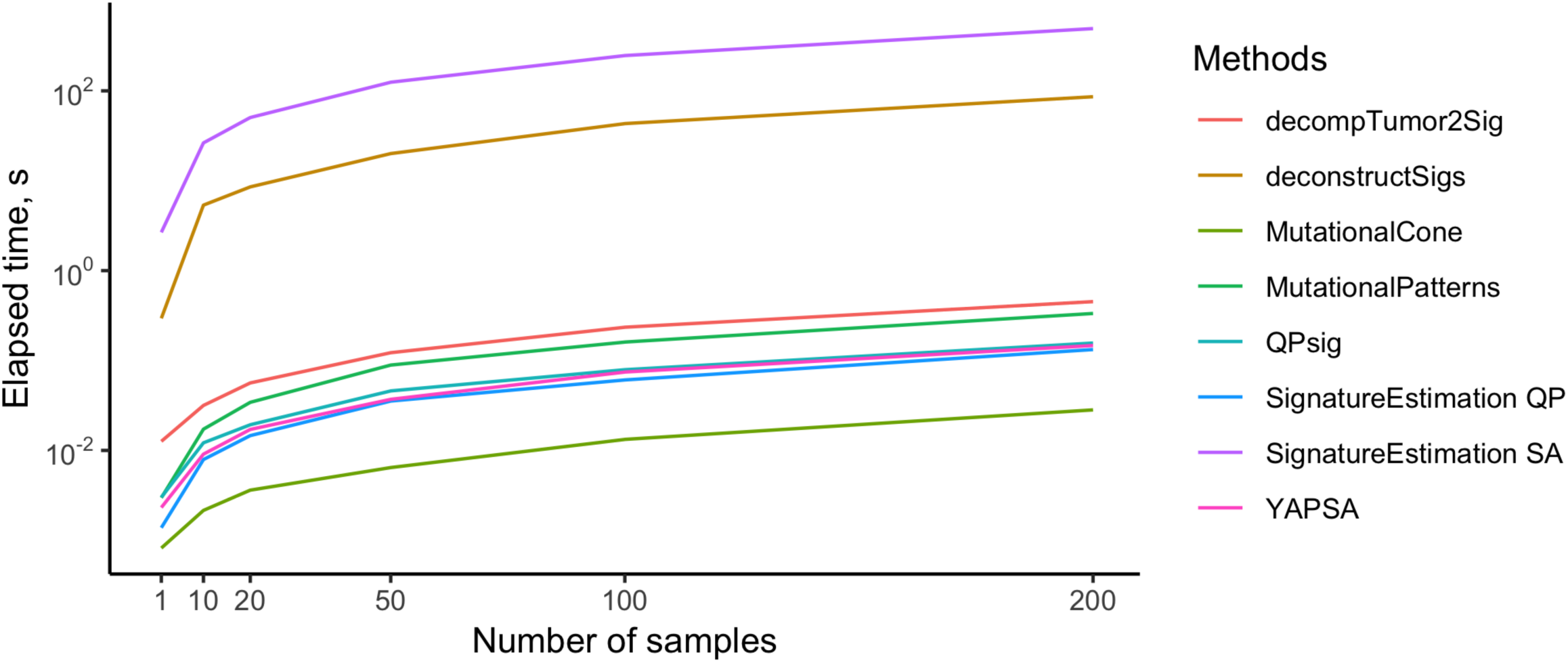
Running times of refitting tools. Methods were applied to subsets of the TCGA Lung cohort of different sizes.

## Discussion

In this work we complement and expand a recent review of the available methods to identify mutational signatures [37] by empirically comparing their performance, using both real world TCGA data and simulated data. To our knowledge, this is the first formal comparison of both de novo discovery and refitting of mutational signatures. The research about mutational signatures is very active and in rapid development, with preprints about new methods regularly published on bioRxiv.org; in this work we thoroughly assessed and formally compared methods that have already been published in peer reviewed journals, and we briefly described other more recent methods. The results of this work can lead to a better understanding of the strengths and limitations of each method as well as to the identification of the key parameters influencing their performance, namely the number of mutations and the “complexity” of the contributing signatures.

Since the publication of the first work about mutational signatures in 2013 [2], multiple algorithms have been developed, leading to similar but not identical results, a source of concern for researchers interested in this type of analysis. Conceptually, this is not surprising: mutational signatures are naturally defined in terms of non-negative matrix factorization, a well-known ill-posed problem (a unique solution does not exist). Although this limitation has cast doubts on the biological validity of mutational signatures, this has been somehow validated using experimental and computational approaches by Zho et al. [27].

An original aspect of this paper is the empirical performance study and in particular the probabilistic model of mutational catalogues used to generate simulations. This model is based on the zero-inflated Poisson distribution that allows for sparse contribution of signatures and thus makes it possible to build mutation count data that are more realistic than the pure Poisson model previously considered [12,13]. As argued in the Simulated data section, our model underrepresents the few samples with extremely large total mutation counts. Because catalogues of this type are more informative for mutational signatures, it is likely that our results slightly underestimate the methods performance. However, our main objective was the comparison of the different methods, and this is not affected by this systematic bias. As discussed, it would have been possible to consider even more realistic models, however these would have led to results that depend on too many parameters thus making the interpretation harder. For the opposite reason, we did not simulate catalogues from real data using the bootstrap: this would have produced almost real samples but without the possibility to evaluate the performance of methods according to different parametric scenarios. Alternative models that were recently proposed are based on the negative binomial distribution [24] and on the Dirichlet distributions for the exposures and signatures and the multinomial distribution for the catalogues [12]. We suggest that developers should assess their new methods on simulations based on realistic models such as ours or the latter.

Another innovative aspect of our work is the assessment of the performance of de novo approaches. This is in general a slippery problem given the non-supervised learning nature of NMF and concurrent methods. We addressed it by building confusion matrices after comparing the “true” signatures used for simulations with the newly identified signatures in terms of their cosine similarity.

At last, we introduced MutationalCone, a new implementation of existing refitting tools that is based on a very simple geometric model and proved to be faster than all other methods. The R code implementing MutationalCone can be found in the Supporting information.

The results of our study can be helpful to guide researchers through the planning of mutational signature analysis. In particular, we show that the performance of de novo methods depends on the complexity of the analysed sequences, the number of mutations and to a lesser degree the number of samples analysed. We showed that cancer profiles for which it is “easier” to detect signatures are characterised by one predominant signature only (as in our profiles 2 and 3), while cancers with more complex profiles, characterized by non-negligible contributions by several signatures (profiles 1 and 4), are more difficult to identify. Indeed, recent evidence shows that the majority of cancers harbour a large number of mutational signatures [23] and therefore belong to the latter scenario.

With regards to the mutation number, we observe that with the number of mutations that could be found in some cancer exomes the performance is generally poor (i.e. low specificity and sensitivity). A clear example is illustrated in a recent comparison between a smaller set of genomes (G=2,780) and a much larger sample of exomes (G=19,184), where it is shown that in the former set a much larger number is signatures is detected than in the latter [23]. This problem is likely to be mitigated if counts were normalized by the expected number of each type’s trinucleotides in the analysed region under healthy condition, that is if an opportunity matrix was provided. We do not address this important aspect in our comparison study as only a few methods can account for opportunity matrices.

We showed that when comparing identified signatures with COSMIC signatures, the choice of a cosine similarity cut-off has a relatively small impact on the overall performance. If the aim is to identify novel signatures it would be preferable to choose a lower value (0.75 or less). On the contrary, if the aim is to access the presence of known signatures in mutational catalogues (cancer genomes or exomes), we recommend turning to refitting methods.

Not all the de novo methods we evaluated offer the possibility to automatically choose the number of signatures to be found. For instance, the popular SomaticSignatures only provides a graphical visualisation of the residual sum of squares for several choices of the number of signatures; the user can choose the optimal number by identifying the inflexion point. For this reason, this crucial aspect was not addressed in our empirical assessment. Similarly, we only considered mutation types defined by the trinucleotide motifs, as currently only pmsignature [18] can consider more than one flanking basis on each side of the substitution.

For well-studied cancers, refitting approaches are a faster and more powerful alternative to de novo methods, even with just one input sample. As the COSMIC database has been built and validated by analysing tens of thousands of sequences of most cancer types, we recommend borrowing strength from previous studies and using refitting tools when performing standard analysis not aimed at the discovery of new signatures.

A more recent set of 49 signatures have been estimated from a new large dataset including thousands of additional sequences. The article introducing this new set has not been published yet and the 30 COSMIC signatures can still be considered as the reference. In the future it will be interesting to evaluate the performance of the different tools using the new set of signatures.

It is important to note that our empirical comparison did not allow to identify a method that is clearly superior to others; for this reason we recommend to systematically perform a sensitivity analysis based on the application of one or more alternative methods based on different algorithms, in order to assess the robustness of results.

## Contributorship

Conceptualization and supervision: GS and VP; formal analysis: HO; writing (original draft preparation, review and editing): HO, GS and VP.

## Acknowledgements

We acknowledge the support of a grant from La Ligue contre le Cancer (Appel à projets 2016 ‘Recherche en Épidémiologie’). HO is supported by a PhD fellowship from the French Institut National du Cancer (INCa_11330). VP would like to thank Grégory Nuel for the useful discussions around zero-inflated models. The authors would like to thank an anonymous reviewer for the very useful remarks and constructive suggestions.

## Supporting information

**S1 Fig.**
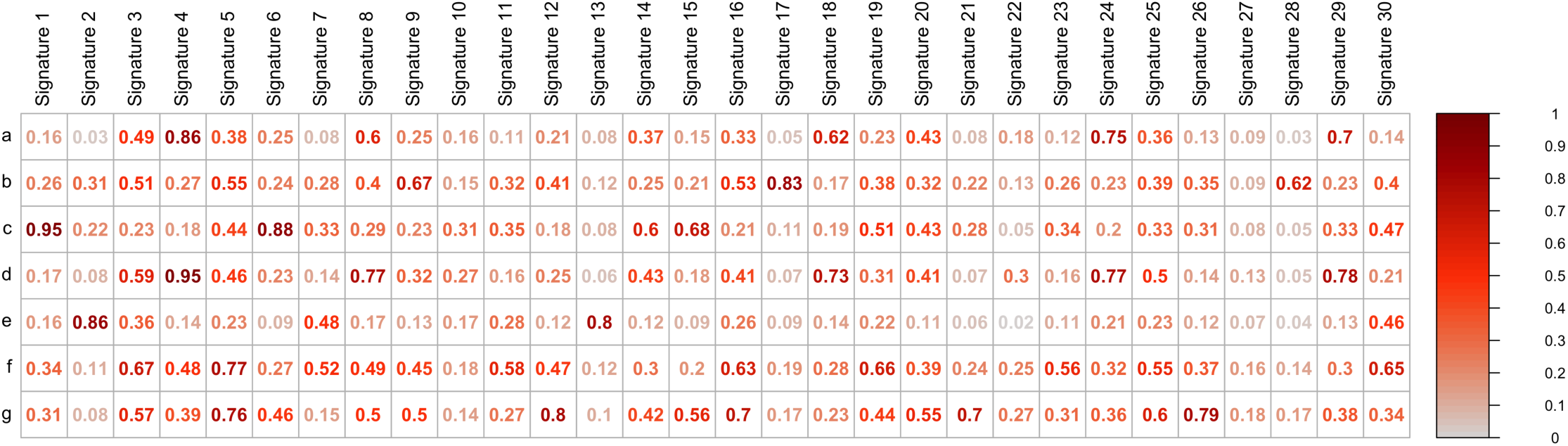
Comparison of newly identified signatures with COSMIC signatures. Signatures a-g were identified in a de novo extraction using the maftools R package from the TCGA Lung Adenocarcinoma cohort of 563 cancer genomes. The novel signatures were then compared to the 30 signatures validated in the COSMIC database in terms of cosine similarity. Each signature is then assigned to the most similar COSMIC signature provided that their cosine similarity is above a fixed threshold. For instance, signature f is matched to signature 5 at a cut-off of 0.75 but is considered as a completely new signature if the cut-off is at 0.80. Also note that a unique assignment can be controversial: for instance, signature g is similar both to signatures 12 and 26.

**S2 Fig.**
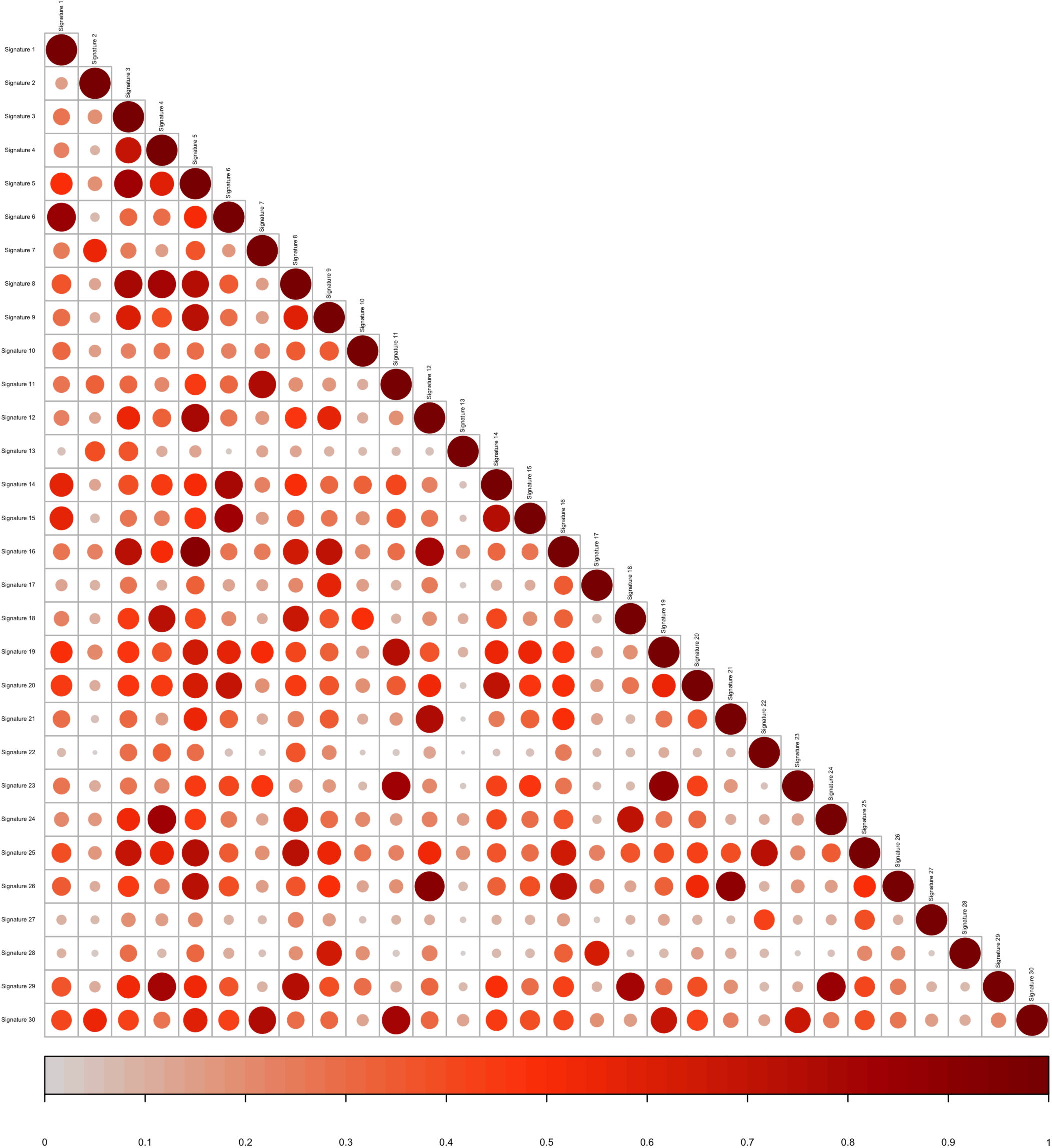
Cosine similarity plot of COSMIC signatures. Some COSMIC signatures are very similar to others. For instance, signature 8 is similar to signatures 3, 4 and 5.

**S3 Fig.**
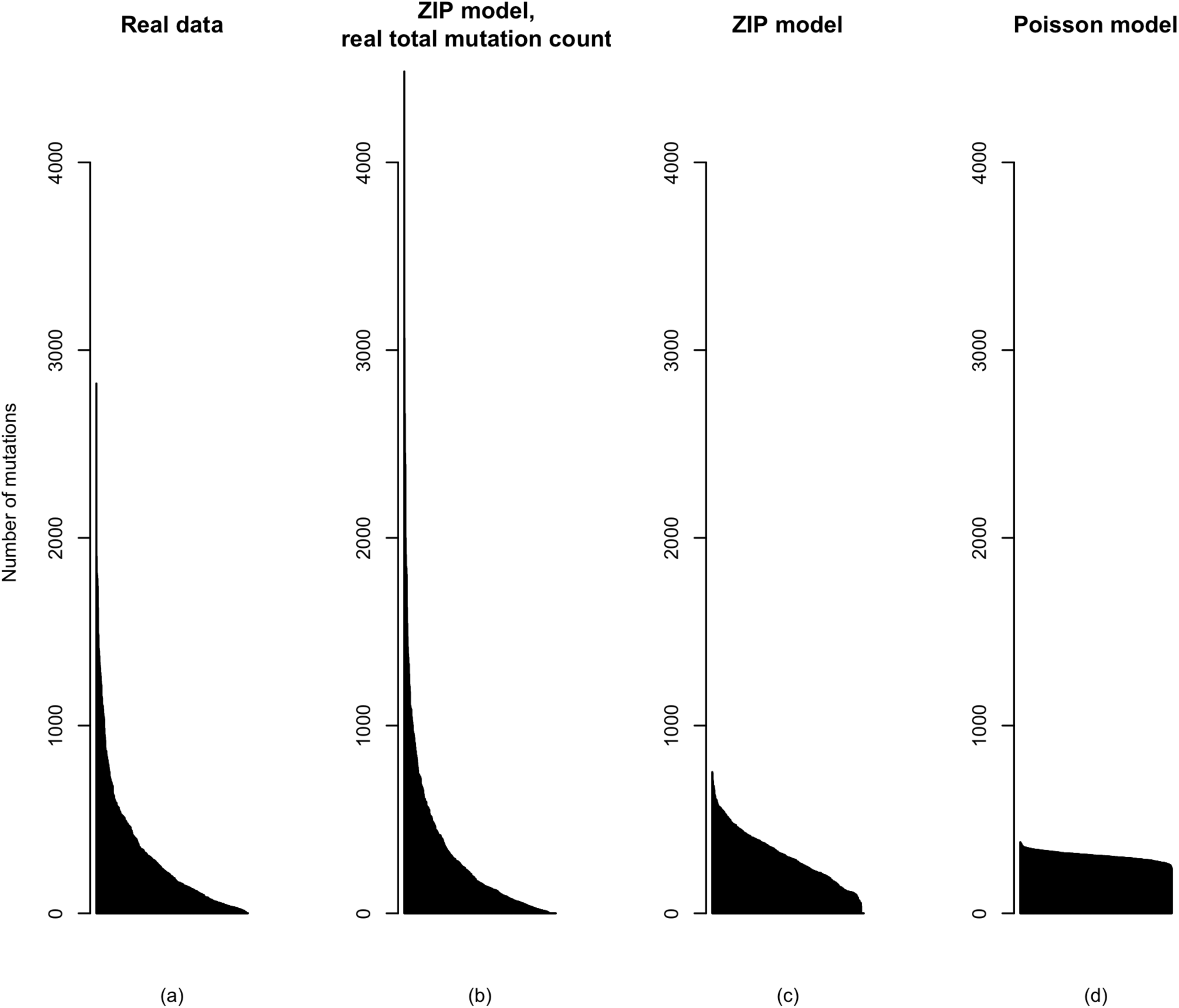
Simulations of 563 lung adenocarcinoma catalogues according to different models. **(a)** Real catalogues from the TCGA lung adenocarcinoma cohort. **(b)-(c)** Catalogues sampled from the ZIP model described in the main text. The relative contribution *q*_*n*_ of each signature *n* is the mean of the relative contributions of *n* in all samples as estimated by Maftools. In **(b)** simulated and real catalogues are in a 1 to 1 correspondence: for each simulated sample *g*, the total number of mutations *r*_*g*_ in the corresponding real catalogue is taken. In **(c)** all samples are simulated according to *r* = 306, the average total number of mutations in the real data. The latter example illustrates the parametric model used for the main empirical study. **(d)** Catalogues sampled according to the Poisson model 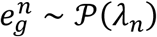, where *λ*_*n*_ is the mean number of mutations due do to *n* in the real samples as estimated by Maftools.

**S4 Fig.**
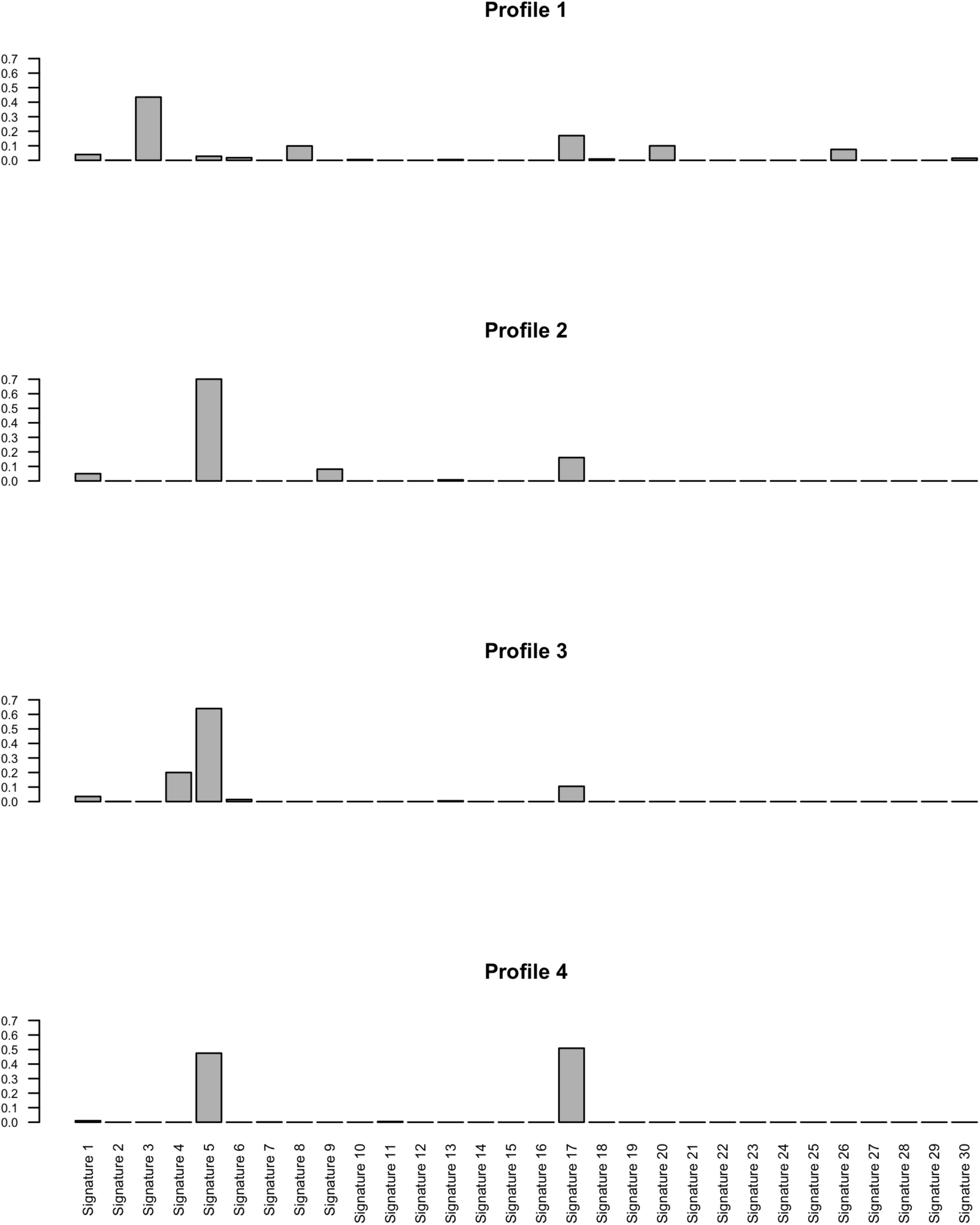
Choice of parameters *q*_*n*_ in the simulations. Four different configurations (*q*_1_,…, *q*_30_) were considered for simulating realistic data. Each configuration represents the average share of mutations due to the different COSMIC signatures and was chosen to mimic real exposure profiles for four cancer types: estimates were obtained from Breast Cancer (Profile 1), Lymphoma (Profile 2), Lung Adenocarcinoma (Profile 3) and Melanoma (Profile 4) TCGA cohorts.

**S1 File. Mutational Cone.** Description of our original method for signature refitting and the corresponding R script.

## MutationalCone

We report here the R code implementing our original method for signature refitting.

Let be the linear subspace of spanned by the reference signatures. Our function MutationalCone() projects the input mutational catalogue onto the cone in spanned by the reference signatures with the very fast coneproj R package (https://cran.r-project.org/web/packages/coneproj). Because projections are simply calculated as scalar products, this function requires the user to specify an orthonormal basis of toghether with the components of the reference signatures with respect to it. These two input matrices can be calculated with the function SignatureSubspace() once and for all, before iterating MutationalCone() on all catalogues. SignatureSubspace() finds an orthonormal basis of with the Gram-Schmidt algorithm.

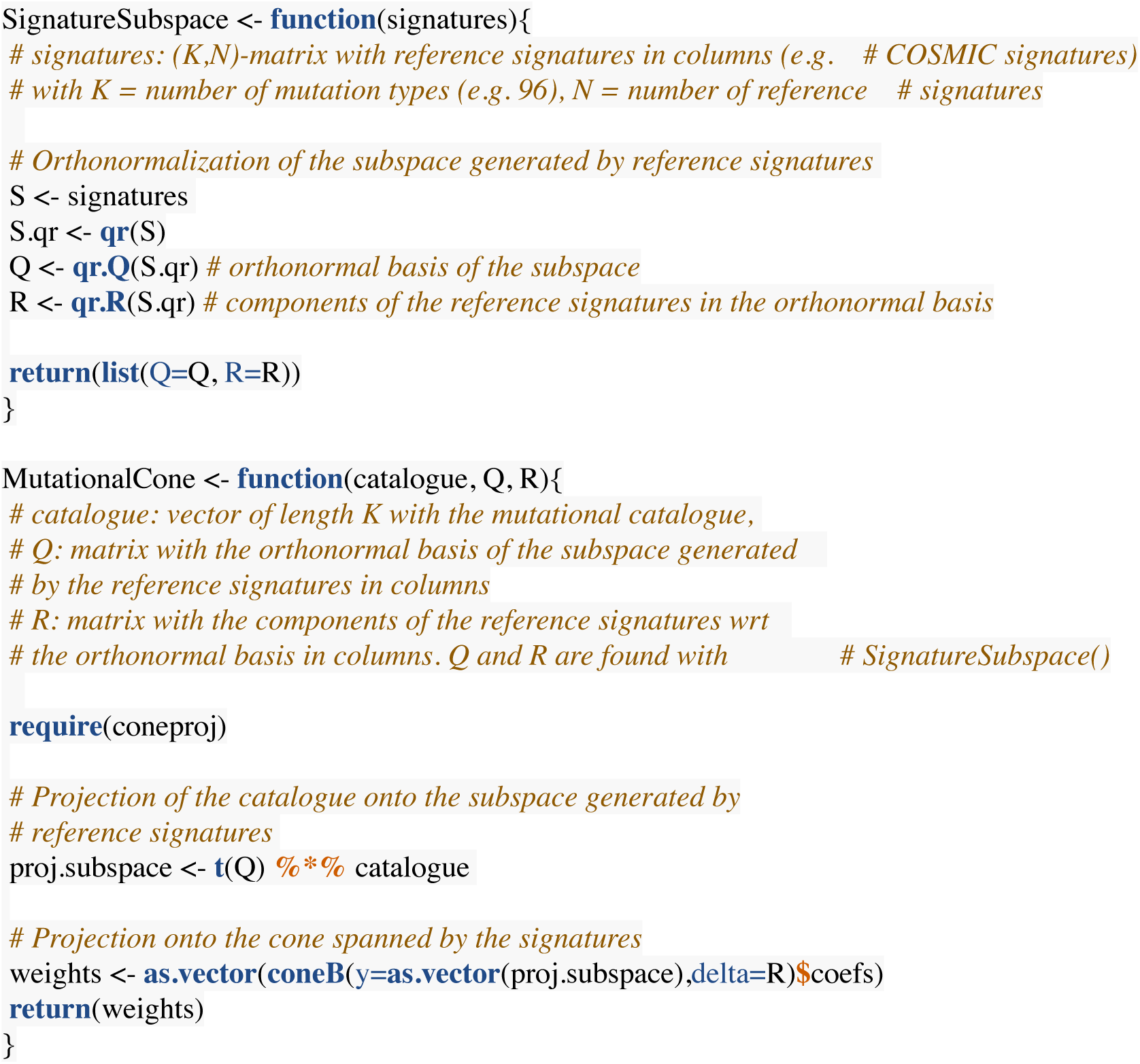

## Notes

#### Summary of Updates

This revised version reviews additional computational tools for mutational signatures that have been recently published.

